# Prevalence of the parasitic copepod, *Sarcotaces* sp., infection in British Columbia rockfishes (*Sebastes* sp.) and implications for rockfish life-history

**DOI:** 10.1101/2025.08.07.668991

**Authors:** Matthew R. Siegle, Jillian C. Dunic, Kristina Castle, Malcolm Wyeth, Milton S. Love, Sean C. Anderson

## Abstract

Marine fishes often suffer detrimental effects of parasitism, which can affect multiple life-history traits, such as maturity, growth, fecundity, and mortality. Rockfishes (genus *Sebastes*), in the northeast Pacific Ocean, are commonly parasitized by copepods in the genus *Sarcotaces*. These copepods form large cysts within the body cavity, or in the musculature of individuals. There is a lack of baseline data on the prevalence of *Sarcotaces* sp. infection and the effect of infection on rockfish health and life-history. Here, we present data on *Sarcotaces* sp. infection in 23 rockfish species (including two thornyhead species, genus *Sebastolobus*) that includes over 37,000 individual records collected on Fisheries and Oceans Canada’s (Pacific Region) groundfish research surveys. *Sarcotaces* sp. were found in all geographic areas covered by the surveys (the coastal waters of British Columbia), and at all depths fish were encountered. Infection rates in the various species ranged from 0% to 10%, and were highest for Pacific Ocean Perch (POP), Silvergray, Rougheye/Blackspotted Complex (REBS), Yellowmouth, and Yelloweye rockfishes. We found that *Sarcotaces* sp. infection shifted the length-at-maturity ogive, with infected individuals maturing at larger sizes—particularly for males, who exhibited a 22.3 mm increase (95% CrI [credible interval]: 5.5, 39.8) in length-at-maturity across species. For most species, this effect was greater in males than in females, and the effect was most pronounced in POP, Silvergray, and REBS rockfish. Moreover, we found that immature males were 2.5 times more likely to be infected than mature males (95% CrI: 1.3, 4.7) and immature females were 1.7 times more likely to be infected than mature females (95% CrI: 0.9, 3.0). While speculative, this is consistent with *Sarcotaces* sp. infection acting as a co-morbidity, possibly affecting fish health and the probability of perishing before reaching maturity. Body condition was largely unrelated to infection status; however, a slight increase (≈ 5%) in condition was associated with infection in REBS, Yelloweye, and Yellowmouth rockfishes. To our knowledge, this is the first large-scale report on *Sarcotaces* sp. infection within rockfishes and on rockfish life-history traits. Future work is needed to elucidate how infection influences other life-history traits and, importantly, whether these changes affect key stock assessment parameters and, ultimately, catch advice provided to fisheries managers.

## 1 Introduction

Parasites are nearly ubiquitous in marine fishes. Detrimental effects of fish parasites on their hosts include reductions in body condition and growth rate, increased susceptibility to predtion, and higher mortality. Life-history responses to parasitism may include changes in fecundity (Gabagambi et al. 2020) or to the size or age at maturity (Lafferty 1993). The extent to which these life-history responses affect population parameters, however, is not well understood. Furthermore, underestimating the effects of parasites may lower the quality of harvest advice given to management for species targeted by fisheries (Lloret et al. 2012). Improving our understanding of parasite-host dynamics and effects on population parameters will improve our stock assessments and the quality of scientific advice for fisheries managers.

The copepods in the genus *Sarcotaces* (Phylichthyidae) are idiosyncratic parasites, found world-wide in several primarily benthic-associated marine fishes (e.g., Komai 1924, Izawa 1974, Moser et al. 1985, Reimer 1991, Ibrahim and Attia 2023). Members of this genus form large cysts in which live a large, amorphous-shaped female (1–2 cm long), one to several males (approximately 2–3mm long), along with eggs and nauplii (Moser et al. 1985, Bullock et al. 1986, Piasecki et al. 2020). Much to the dismay of biologists and fishers who have accidentally punctured the cysts, the cysts are filled with black fluid, likely derived from blood the parasite has obtained from the host (Kabata 1970). Hosts often harbour multiple cysts (Komai 1924, Moser et al. 1985). In the eastern Pacific, two groups are known to be infected, numerous species of rockfishes (genus *Sebastes*), and Pacific flatnose (*Antimora microlepis* Bean, 1890). While the cysts may be found in various parts of the body (Izawa 1974, Moser 1977, Ibrahim and Attia 2023), in eastern Pacific rockfishes they are typically located in the posterior of the coelomic cavity near the anus, or in the lateral musculature (Kuitunen-Ekbaum 1949, Liston et al. 1960, Moser et al. 1985, Stanley and Kronlund 2005, Conrath 2019, Myers et al. 2019). The genus *Sarcotaces* sp. is classified as a mesoparasite (Piasecki and Boxshall 2024), because, although the majority of the parasite’s body occupies the interior of the host, the female’s last body segment is in contact with the sea. Currently, there are nine recognized (or at least nominal) species in the World Register of Marine Species (WoRMS), although almost no genetic work has been conducted, and the genus is in need of revision (Piasecki et al. 2020). Historically, those *Sarcotaces* sp. found along the eastern Pacific have been referred to as *Sarcotaces arcticus* (Collett 1874). However, because of this taxonomic uncertainty, in this paper we will refer to the parasite as *Sarcotaces* sp..

In general, there is a lack of baseline data on *Sarcotaces* sp. infection in eastern Pacific rock-fishes, and even less is known about rockfish life-history responses to infection. *Sarcotaces* sp. may affect rockfish reproductive success, as it has been implicated in female reproductive failure, through physically blocking sperm from reaching eggs (*S. ciliatus*) (Conrath 2019), possibly through castration (*S*. hopkinsi, M. Love, unpubl. data), and perhaps through reductions in fecundity (suggested for *S. brevispinis*, Stanley and Kronlund 2005). The lack of baseline information on *Sarcotaces* sp. infection rates and effects on life-history responses are particularly worrisome if infection has downstream effects on key stock assessment parameters, such as age and size-at-maturity, fecundity, and selectivity.

Here, we present the first large-scale investigation into the extent of *Sarcotaces* sp. infection in a suite of northeastern Pacific rockfishes across the coastal waters of British Columbia, Canada. We show the spatial extent of *Sarcotaces* sp. encounters, present length-at-maturity ogives for infected and uninfected individuals for a subset of rockfish species, compare differences in infection rates between reproductively mature and immature individuals, and assess potential effects of *Sarcotaces* sp. infection on body condition.

## 2 Methods

### 2.1 Groundfish surveys

Fisheries and Oceans Canada (DFO) conducts a suite of annual fishery-independent surveys using bottom trawl, longline hook, and longline trap gear that, in aggregate, provide comprehensive coverage for the waters of Canada’s Pacific Coast. The core surveys have in common full enumeration of the catches, size composition sampling for most species, and more detailed biological sampling of selected species.

There are four core synoptic bottom trawl surveys, which operate every other year. The Hecate Strait (HS) and Queen Charlotte Sound (QCS) surveys are conducted in odd-numbered years while the West Coast Vancouver Island (WCVI) and West Coast Haida Gwaii (WCHG) surveys are conducted in even-numbered years. All of the synoptic bottom trawl surveys along the British Columbia (BC) coast follow the same random depth-stratified design. Each survey area is divided into 2 km by 2 km blocks and each block is assigned one of four depth strata based on the average bottom depth in the block. The four depth strata vary between areas and range from 50–1300 meters of trawlable habitat. For each survey and in each year, blocks are randomly selected within each depth stratum. Each selected block is inspected and assessed to be rejected as unfishable if one or more unsuccessful attempts are made to fish the block.

In addition to the annual bottom trawl surveys, two area-specific random depth-stratified longline hook surveys known as Hard Bottom Longline Hook (HBLL) Surveys are conducted each year. The commercial longline hook fishing industry contracts vessels and sea-going technicians for a survey of “outside” waters (those occurring outside of the area between Vancouver Island and the mainland) while a separate longline hook survey of “inside” waters (between Vancouver Island and the mainland) is conducted by DFO staff onboard a Canadian Coast Guard research vessel. Both longline surveys occur on a two-year time frame as well, where the northern areas are sampled in odd-numbered years and the southern areas are sampled in even-numbered years. The HBLL surveys also use a 2 km by 2 km random depth-stratified design, where a subset of blocks with one depth stratum per block are randomly sampled each year. The HBLL outside survey samples three depth strata (20–70m, 71–150m, 151–260m), and the HBLL inside survey samples two depth strata (40–70m and 71–100m). These longline surveys provide coast-wide coverage of most of the non-trawlable habitat between 20 and 260 meters depth that is not covered by the bottom trawl surveys.

The final randomized survey conducted each year is a coast-wide longline trap survey targeting Sablefish (*Anoplopoma fimbria*), known as the Sablefish Research and Assessment Survey. The commercial Sablefish fishing industry supplies a chartered commercial fishing vessel and the survey is conducted with a combination of DFO staff and industry-hired sea-going technicians. The stratified random portion of the survey covers the depth range of 180–1370 m for the entire outer BC coast, while a fixed-station portion covers several central coast inlets.

In addition to these core annual surveys, we include data from other research surveys that are used within some groundfish stock assessments. These surveys include the WCVI Multispecies Small-mesh Bottom Trawl Survey, the Strait of Georgia Dogfish Survey, the International Pacific Halibut Fishery-Independent Setline Survey, and the joint Canada/U.S. Hake Acoustic Survey.

### 2.2 Biological sampling

In addition to the full enumeration of the catches and size composition sampling for most species, more detailed biological sampling of selected species are conducted on all of the DFO groundfish surveys. Most rockfishes are among the species selected for more detailed biological sampling. The biological sampling for rockfishes includes: individual length, individual weight, sex, and maturity status (macroscopic identification, Table S1). Otoliths are taken for subsequent ageing by the DFO Sclerochronology Lab (however, only a subset of individuals are selected for age analysis), and a fin clip is taken from a subset of rockfishes for DNA analysis by various agencies, including DFO. The HBLL inside survey also collects stomach content data from the sampled rockfishes and Lingcod (*Ophiodon elongatus*).

*Sarcotaces* sp. presence within rockfishes was sporadically recorded beginning in 1969. These earlier records include presence-only information, and it is unclear if null records correspond to no data or the absence of *Sarcotaces* sp.. We therefore excluded these samples from our analyses. Systematic sampling for *Sarcotaces* sp. infection began as a standard part of the biological sampling in 2019 and provides true presence/absence and count data.

### 2.3 Statistical methods

Here, we provide an overview of the statistical methods. A full description of the statistical methods with a complete list of equations is provided in the Supporting Information.

#### 2.3.1 Spatial modelling of *Sarcotaces* sp. infection probability

We modelled the spatial probability of *Sarcotaces* sp. encounter with a logistic regression with a quadratic effect for log depth, an independent mean for each year, and a Gaussian Markov random field (GMRF) to account for latent spatial factors driving spatial correlation.

We estimated anisotropy in the GMRF (Haskard 2007, Fuglstad et al. 2015) to allow for spatial correlation to decay differently up and down the coast from off the coast and fit the model using a finite-element mesh (Lindgren et al. 2011, Lindgren 2025) with a minimum triangle edge length of 5 km. We fit this spatial model with the sdmTMB R package (Anderson et al. 2024), which uses fmesher (Lindgren 2025) to generate input matrices and TMB (Kristensen et al. 2016) to calculate the marginal likelihood.

#### 2.3.2 Bayesian modelling of length- and age-at-maturity

We estimated length-at-maturity with hierarchical Bayesian logistic regression models fit separately to male and female fish:

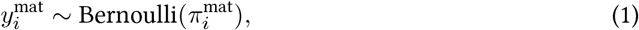

where the probability of maturity, 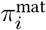, is modeled as:

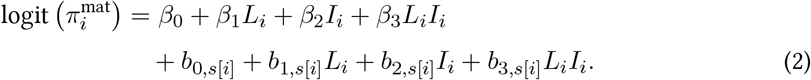

Here, 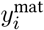 is either mature (1) or immature (0) for fish *i*; *β*_0_, *β*_1_, *β*_2_, and *β*_3_ are fixed-effect coefficients; *L*_*i*_ is the (standardized) length of fish *i*; *I*_*i*_ is the presence (1) or absence (0) of *Sarcotaces* sp. infection in fish *i*, and *b*_0,*s*[*i*]_, *b*_1,*s*[*i*]_, *b*_2,*s*[*i*]_, and *b*_3,*s*[*i*]_ are species-specific random intercepts and slopes for species *s*, fish *i*.

Since there were fewer data with ages available (total n = 2,334, infected n = 37), we were unable to fit the above hierarchical model to estimate age-at-maturity. Instead, we fit a simplified model of the probability of age-at-maturity as:

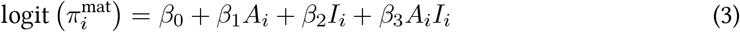

where symbols are as described above with the addition of *A*_*i*_ representing standardized age of fish *i*. We used the same priors as in Equations S7 and S8 (see Supporting Information).

To compare *Sarcotaces* sp. infection probability between fishes of different sex and maturity, we fit additional hierarchical logistic regressions. We first compared infection rates between immature and mature individuals, and then compared differences across maturity stages. Given the relatively low sample sizes of individuals in maturity stages 4 and 5 (Table S1), we combined these stages for this analysis. We also tested for differences in mean age between infected and uninfected individuals (pooled across species) with the Wilcoxon test (Wilcoxon 1945).

#### 2.3.3 Bayesian modelling of body condition

To evaluate whether *Sarcotaces* sp. infection is associated with changes in body condition, we fit hierarchical linear models to Le Cren’s body condition index (Le Cren 1951). First, for each species and sex, we fit linear regressions of log(weight) against log(length) and removed outliers with residuals that were ≥ 10 SD. We then refit the regressions without the outliers to estimate the standard weight-length relationship: 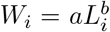, where *W*_*i*_ and *L*_*i*_ are the weight and length of fish *i*. We then used the estimated values of *a* and *b* to calculate Le Cren’s condition index *K*_*i*_ for fish *i* by dividing the observed weight by the expected weight: 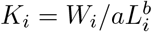.

For each sex, we then fit a hierarchical model of the form:

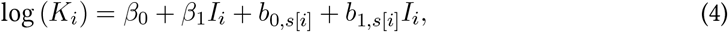

where *K*_*i*_ is the condition index for fish *i, β*_0_ and *β*_1_ are fixed effect coefficients, *I*_*i*_ is the presence (1) or absence (0) of *Sarcotaces* sp., and *b*_0,*s*[*i*]_ and *b*_1,*s*[*i*]_ are species-specific random intercepts and slopes for species *s*, fish *i*.

#### 2.3.4 Bayesian sampling

We fit our Baysian models with the brms R package (Bürkner 2017) using the cmdstanr (Gabry et al. 2025) and Stan (Carpenter et al. 2017) interface to the No-U-Turn Hamiltonian Markov Chain Monte Carlo (MCMC) sampling algorithm (Hoffman and Gelman 2014). For each model, we ran 2000 MCMC iterations on each of four chains, discarding the first 1000 samples of each chain as warm-up. We assessed that our MCMC chains were consistent with convergence through traceplots and by checking that the 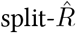 (potential scale reduction factor) statistic was *<* 1.01 and that bulk and tail effective sample sizes were *>* 400 for all parameters (Vehtari et al. 2021). We performed all analyses in R version 4.5.0 (R Core Team 2025).

## 3 Results

We examined over 37,000 individual records from 23 rockfish species (including Longspine and Shortspine Thornyheads [*Sebastolobus altivelis* and *S. alascanus*, respectively]) for *Sarcotaces* sp. count per individual. *Sarcotaces* sp. have been encountered in all geographic areas covered by DFO’s groundfish surveys (Figure 1a). Some rockfish species are encountered in our surveys, but are not prioritized for detailed biological sampling, including sampling for *Sarcotaces* sp. presence (e.g., Harlequin and Pygmy Rockfishes, *S. variagatus and S. wilsoni*). These species have sample sizes fewer than 50 individuals in this dataset. Other species are frequently encountered (e.g., POP, Quillback, and Yelloweye Rockfishes, *S. alutus, S. maliger, and S. ruberrimus*, respectively), where we have over 5,000 records per species. The encounter rate of *Sarcotaces* sp. per species varies from zero percent to a high of 10% in Yelloweye Rockfish (Table 1). The probability of *Sarcotaces* sp. encounter is positive in all areas of the BC coast; however, there is a notable lack of *Sarcotaces* sp. found off the west coast of Haida Gwaii (Figure 1b).

**Table 1:** Sample sizes and infection detections for species examined for *Sarcotaces* sp. infection in 2019–2022. Species highlighted in grey had no observed infections in any specimens examined. Species marked with an asterisk have been found with *Sarcotaces* sp. cysts in non-systematic sampling events.

| Species | Uninfected | Infected | Total n | Infection rate (%) |
| --- | --- | --- | --- | --- |
| Yelloweye Rockfish | 5834 | 647 | 6481 | 10 |
| Rougheye/Blackspotted | 1849 | 152 | 2001 | 7.6 |
| Yellowmouth Rockfish | 911 | 71 | 982 | 7.2 |
| Silvergray Rockfish | 2744 | 132 | 2876 | 4.6 |
| Pacific Ocean Perch | 5321 | 161 | 5482 | 2.9 |
| Redbanded Rockfish | 3385 | 16 | 3401 | 0.5 |
| Widow Rockfish | 208 | 1 | 209 | 0.5 |
| Quillback Rockfish | 5658 | 19 | 5677 | 0.3 |
| Redstripe Rockfish | 1807 | 6 | 1813 | 0.3 |
| Canary Rockfish | 1186 | 3 | 1189 | 0.3 |
| Copper Rockfish | 857 | 3 | 860 | 0.3 |
| Bocaccio | 2188 | 2 | 2190 | 0.1 |
| Shortspine Thornyhead | 1202 | 0 | 1202 | 0 |
| Longspine Thornyhead | 592 | 0 | 592 | 0 |
| Yellowtail Rockfish | 488 | 0 | 488 | 0 |
| Greenstriped Rockfish* | 304 | 0 | 304 | 0 |
| China Rockfish* | 292 | 0 | 292 | 0 |
| Shortbelly Rockfish | 175 | 0 | 175 | 0 |
| Shortraker Rockfish* | 144 | 0 | 144 | 0 |
| Tiger Rockfish | 70 | 0 | 70 | 0 |
| Harlequin Rockfish* | 27 | 0 | 27 | 0 |
| Pygmy Rockfish | 27 | 0 | 27 | 0 |
| Black Rockfish | 12 | 0 | 12 | 0 |

**Figure 1.**
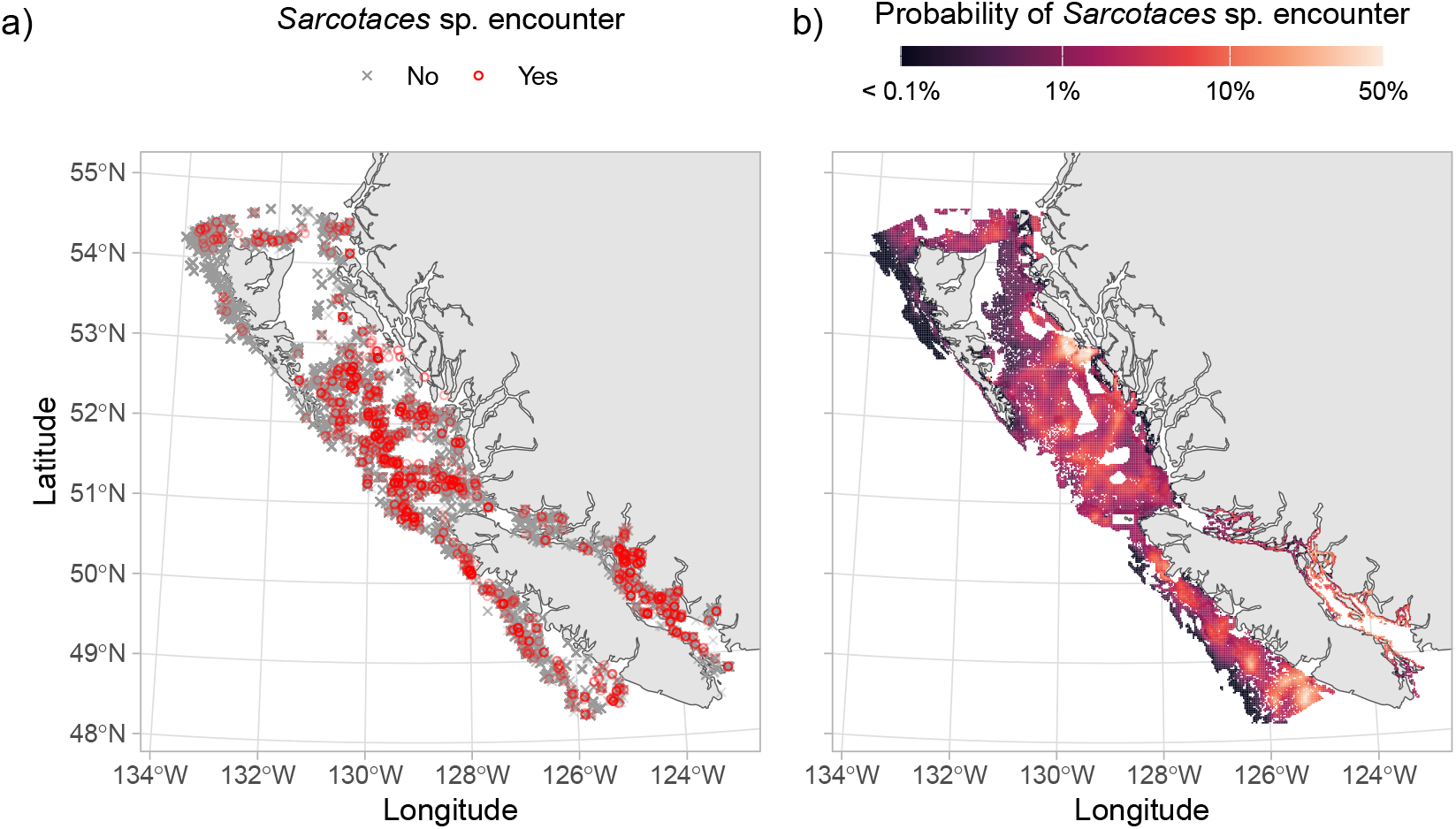
*Sarcotaces* sp. encounters in Fisheries and Oceans Canada’s Synoptic Trawl and Hardbottom longline groundfish surveys. (a) Individual observations. (b) Probability of *Sarcotaces* sp. encounter from a generalized linear model with spatial random effects (Supporting Information).

To date, six species have not been recorded with *Sarcotaces* sp. cysts (Table 1). Of these, Shortbelly Rockfish (*S. jordani*), Tiger Rockfish (*S. nigrocinctus*), Pygmy Rockfish, and Black Rockfish (*S. melanops*) have few samples (*<* 175). Conversely, Shortspine Thornyhead (n=1,202) and Longspine Thornyhead (n=592) were highly sampled with no observed infections. Five species that were systematically sampled between 2019–2022 (grey in Table 1) were not observed with infections; but, in the opportunistic sampling since 1962, infected individuals have been found: Shortraker Rockfish (*S. borealis*, n=5), Harlequin Rockfish (n=4), Greenstriped Rockfish (*S. elongatus*, n=1), and China Rockfish (*S. nebulosus*, n=1).

Most individuals infected by *Sarcotaces* sp. were only parasitized with one cyst. Fewer individuals had two or 3+ cysts, and a handful of individuals had 6+ cysts (Figure 2, Table S2). *Sarcotaces* sp. presence does not appear to be structured by depth. For rockfish species that exhibit rates of infection greater than 2%, *Sarcotaces* sp. are found across the depth range we have encountered for species in our surveys (Figure 3).

**Figure 2.**
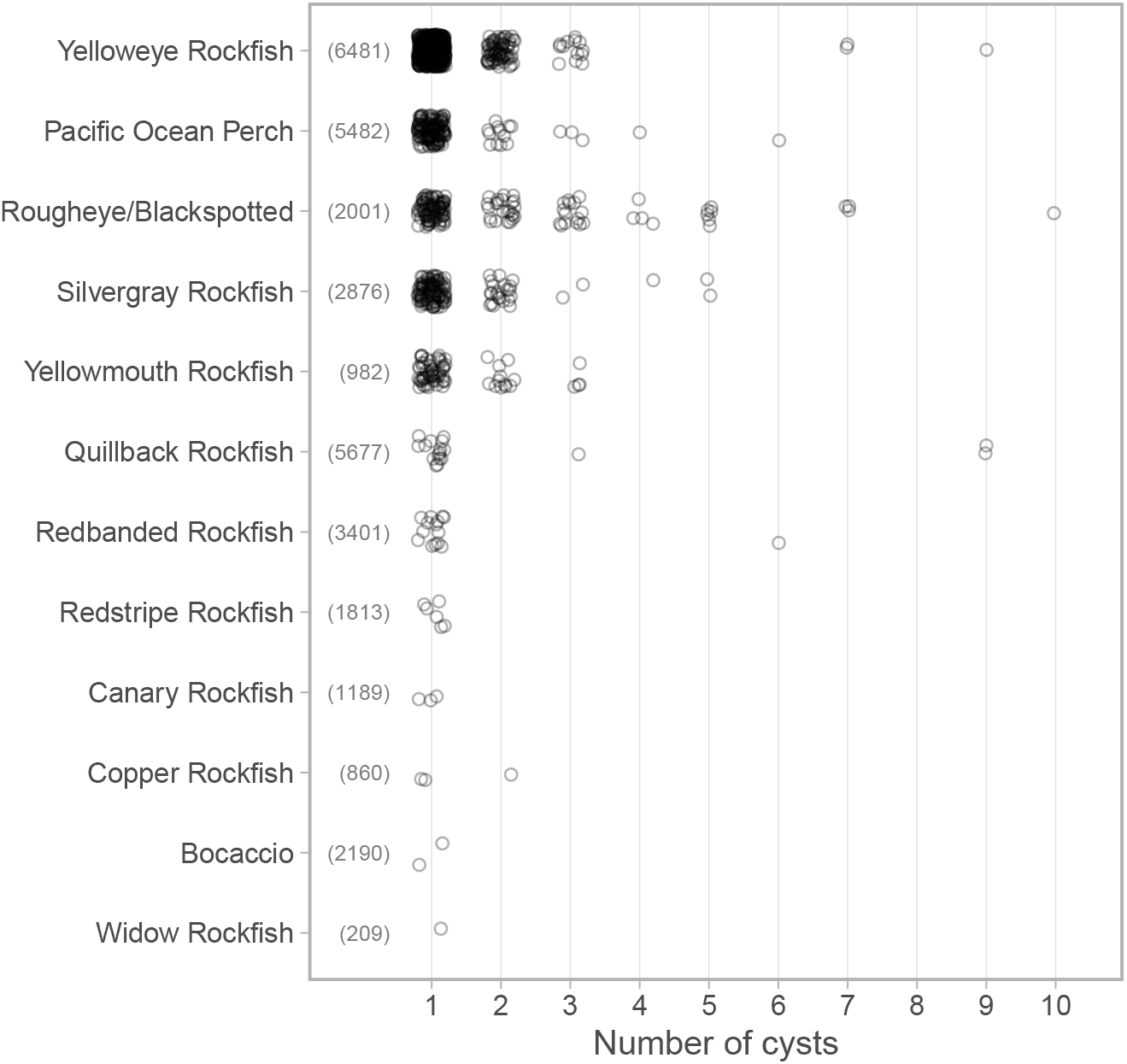
Frequency of *Sarcotaces* sp. cyst counts per individual by species for all fish with at least one *Sarcotaces* sp. cyst. Points have been randomly jittered for visualization. Numbers in parentheses give the total number of systematically sampled fish.

**Figure 3.**
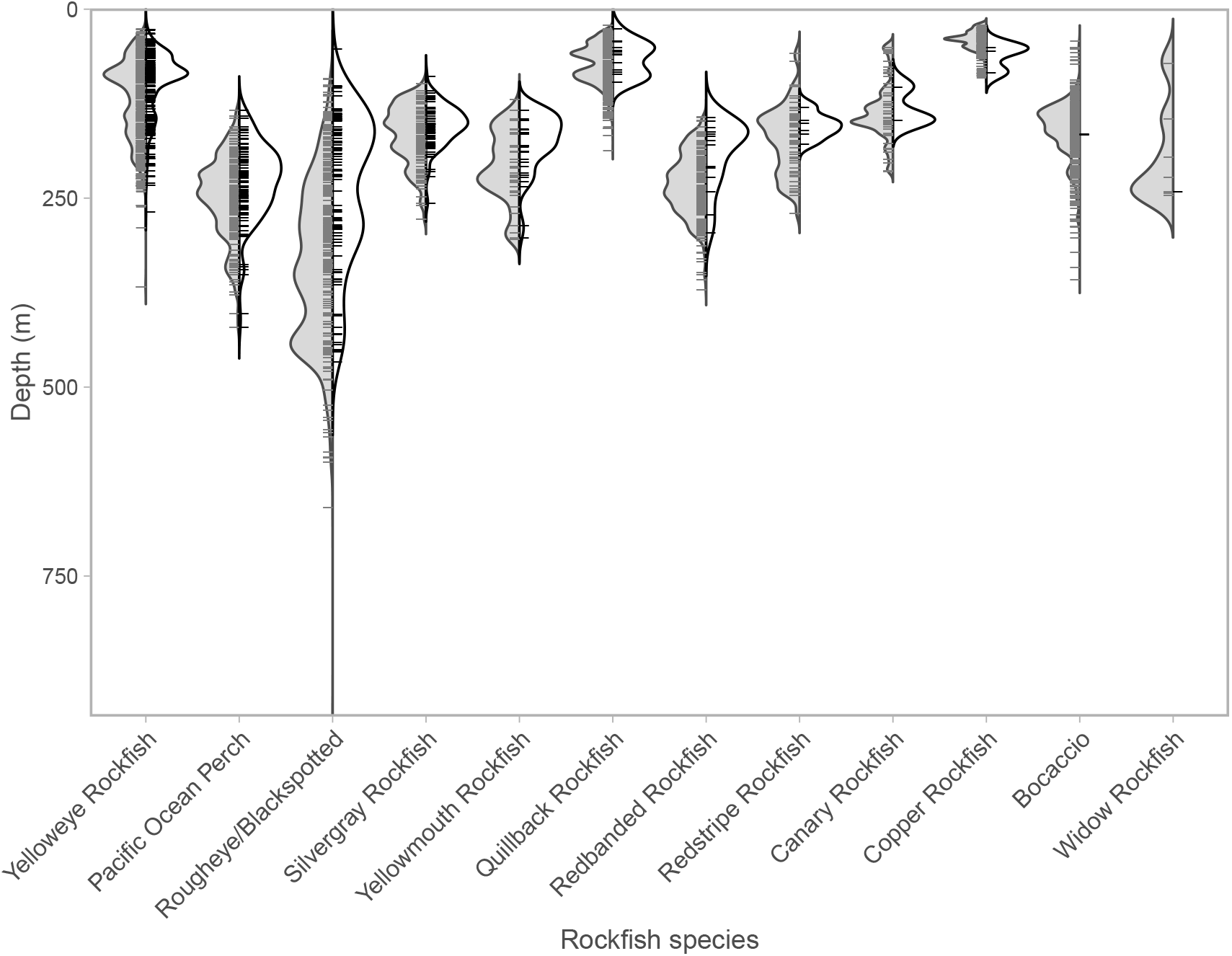
Depth distribution of *Sarcotaces* sp. observations within species. Species are arranged from left to right based on the absolute number of infected individuals. The estimated density of uninfected (grey) and infected (white) individuals is displayed, excluding cases with fewer than two observations. Short horizontal lines represent individual observations.

We observed an effect of *Sarcotaces* sp. infection on the length-at-maturity ogives for both males and females; however, the effect was more pronounced in males (Figure 4a, Figure 5). Across species, the length at 50% maturity was 22.3 (95% CrI [credible interval]: 5.5, 39.8) mm higher for infected individuals than for uninfected. Infected female POP exhibited a length at 50% maturity 7.5 (95% CrI: -7.2, 22.8) mm higher than their uninfected counterparts (Figure 5). Within species, this effect was most pronounced in males of POP, Silvergray, and REBS (Figures 5, S1, S2). For males and females of these three species, the length at 50% maturity was 20–30 mm higher for infected individuals than for uninfected (Figure 5).

**Figure 4.**
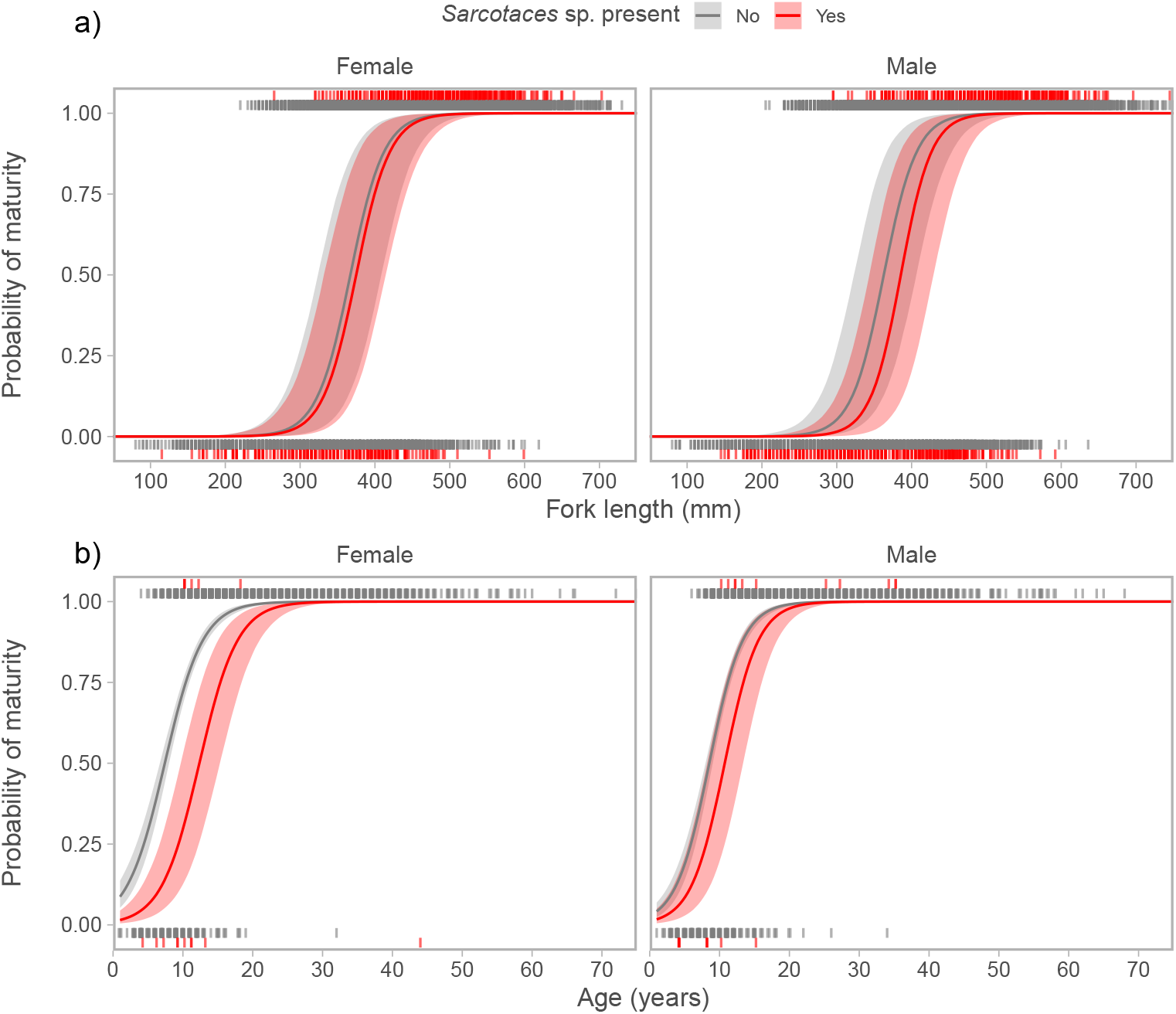
Population-level expectations of rockfish maturity as a function of a) length and b) age in relation to *Sarcotaces* sp. infection, based on fixed effects from a hierarchical model. Species-level effects were excluded to represent the mean effect of *Sarcotaces* sp. infection on maturity at length.

**Figure 5.**
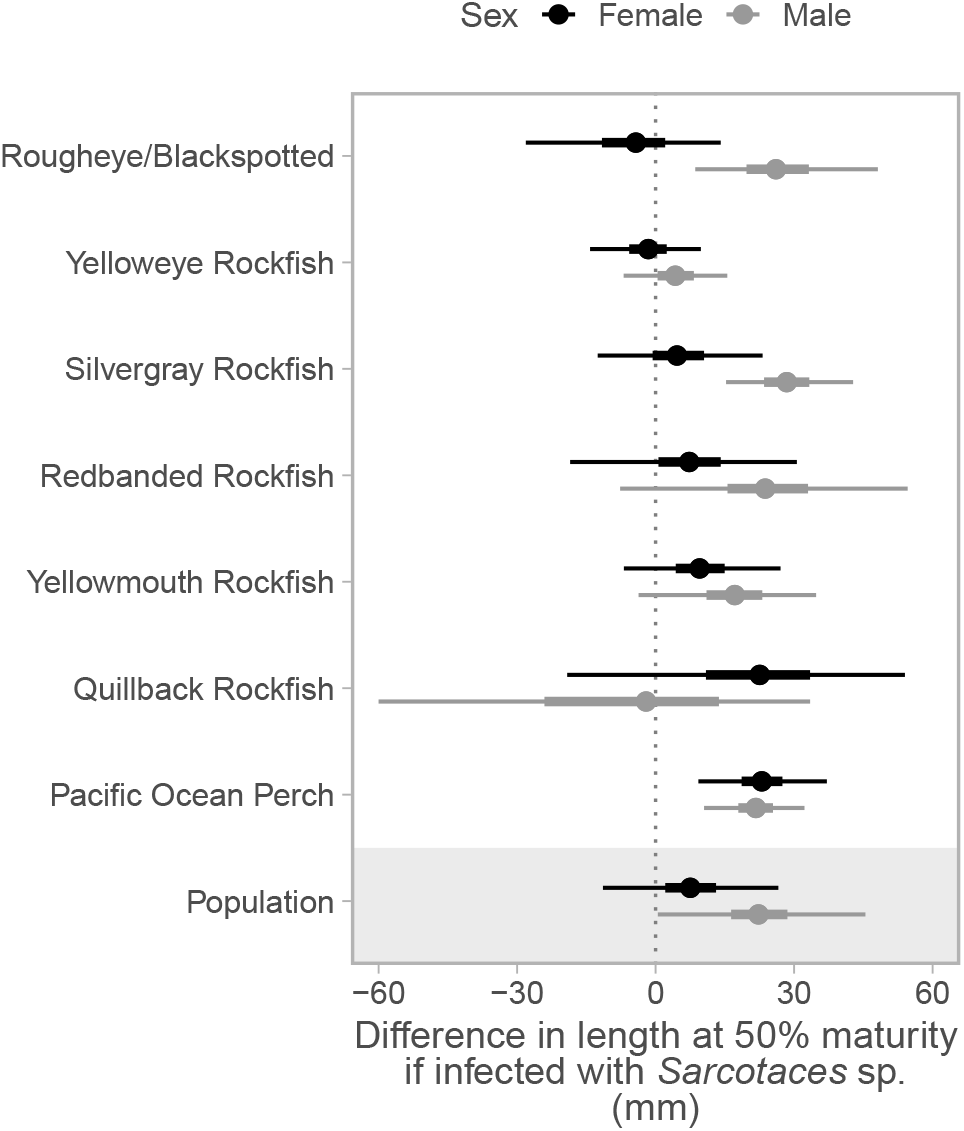
Posterior distribution showing difference in length at 50% maturity for individuals with *Sarcotaces* sp. infection relative to uninfected individuals. Points are posterior medians and thick and thin lines show 50% and 95% credible intervals, respectively. Species are ordered from top to bottom by increasing difference in female length at 50% maturity. The population-level estimate— across all species—is shown at the bottom and reflects the statistical population mean.

Due to limited age data (we only included 37 infected individuals with ages), only three species were included in the age-at-maturity analysis (Pacific Ocean Perch, Quillback Rockfish, and Yellowmouth Rockfish). We observed an overall negative effect of *Sarcotaces* sp. infection on the age-at-maturity for females, but only a weak negative effect in males (Figure 4b). In females, infected individuals reached maturity 5.1 (95% CrI: 2.3, 8.2) years later than uninfected individuals, whereas infected males were 2.2 (95% CrI: -0.4, 4.8) years older on average.

Both reproductively immature males and females exhibited higher rates of *Sarcotaces* sp. infection than their reproductively mature counterparts. We found that immature males had a 74% probability of having two times the infection rate than mature males, and that immature females had a 93% probability of higher infection rates than mature females (Figure 6). Across species, Silvergray, REBS, POP, and Yellowmouth rockfish had the greatest differences, with Yellowmouth males having infection rates in immature males 6.5 (95% CrI: 3.9, 11.5) times greater than mature. Comparison of infection rates by maturity status showed that rates were greatest in status 1 and 2 (immature and maturing) individuals in most species and were similar for maturity status groups 3 to 7 (developing through resting) (Figures S3, S4, S5, Table S1). Infected individuals (pooled across species) also had a lower mean age than uninfected individuals (females: 7.8 [95% CrI: 4.5, 10.3] years lower; males: 5.6 [95% CrI: 2.3, 8.1]] years lower (Figure **??**). However, we caution against over-interpreting this result as the sample size for the infected group was low (n = 37).

**Figure 6.**
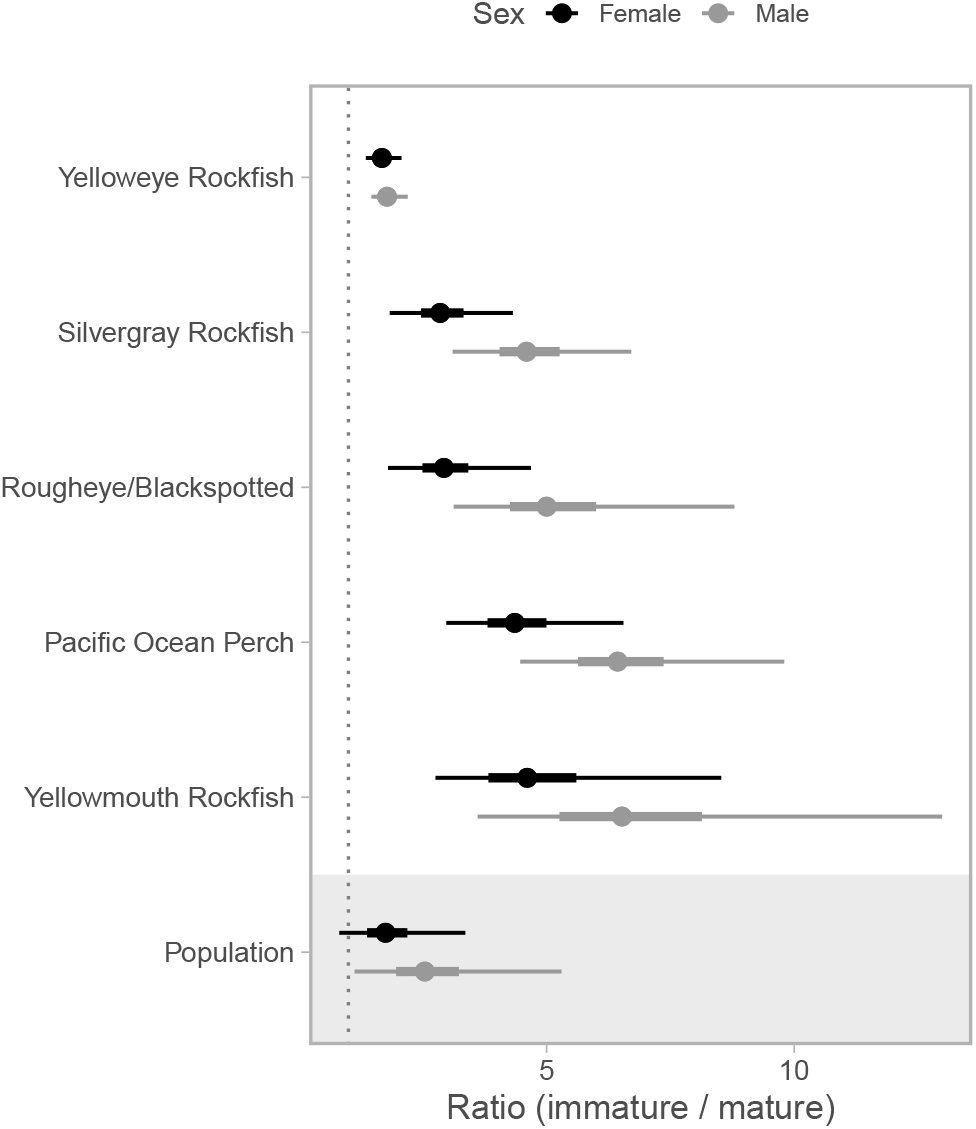
Posterior distribution of the ratio of infection probabilities in immature and mature rockfish. The ratio is calculated as Pr(infection|immature) / Pr(infection|mature). Values greater than 1 (vertical dashed line) indicate higher infection probability in immature individuals. Points are posterior medians and thick and thin lines show 50% and 95% credible intervals, respectively. Species are ordered from top to bottom by increasing ratio of infection rate, with the population-level difference shown bottom. At the population-level there is a 93% probability that immature females have higher infection rates than mature females and a 99% probability that immature males have higher infection rates than mature males. Additionally, there is a 74% probability that immature males have at least twice the infection rate of mature males). Only species with at least 10 infections found in each of immature and mature categories are shown.

Across species, *Sarcotaces* sp. infection had a weak but positive effect on body condition (Figure 7). The probability that an infected individual had higher body condition than uninfected individual was 95% for females and 91% for males. However, the effect sizes were small with high uncertainty; the median estimated increase in body condition was 2.3% in females (95% CrI: -0.1, 4.4) and 1.4% in males (95% CrI: -0.5, 3.1). Across most species, *Sarcotaces* sp. infection was not associated with a difference in body condition (Figure 7). However, in REBS, Yelloweye, and female Yellowmouth, higher body conditions were associated with infection. This effect, represented as the percent increase in body condition given infection, was strongest in both female (5.1%; 95% CrI: 3.2, 7.1) and male (5.0%; 95% CrI: 3.4, 6.8) REBS, and female Yellowmouth (5.0%; 95% CrI: 2.6, 7.7).

**Figure 7.**
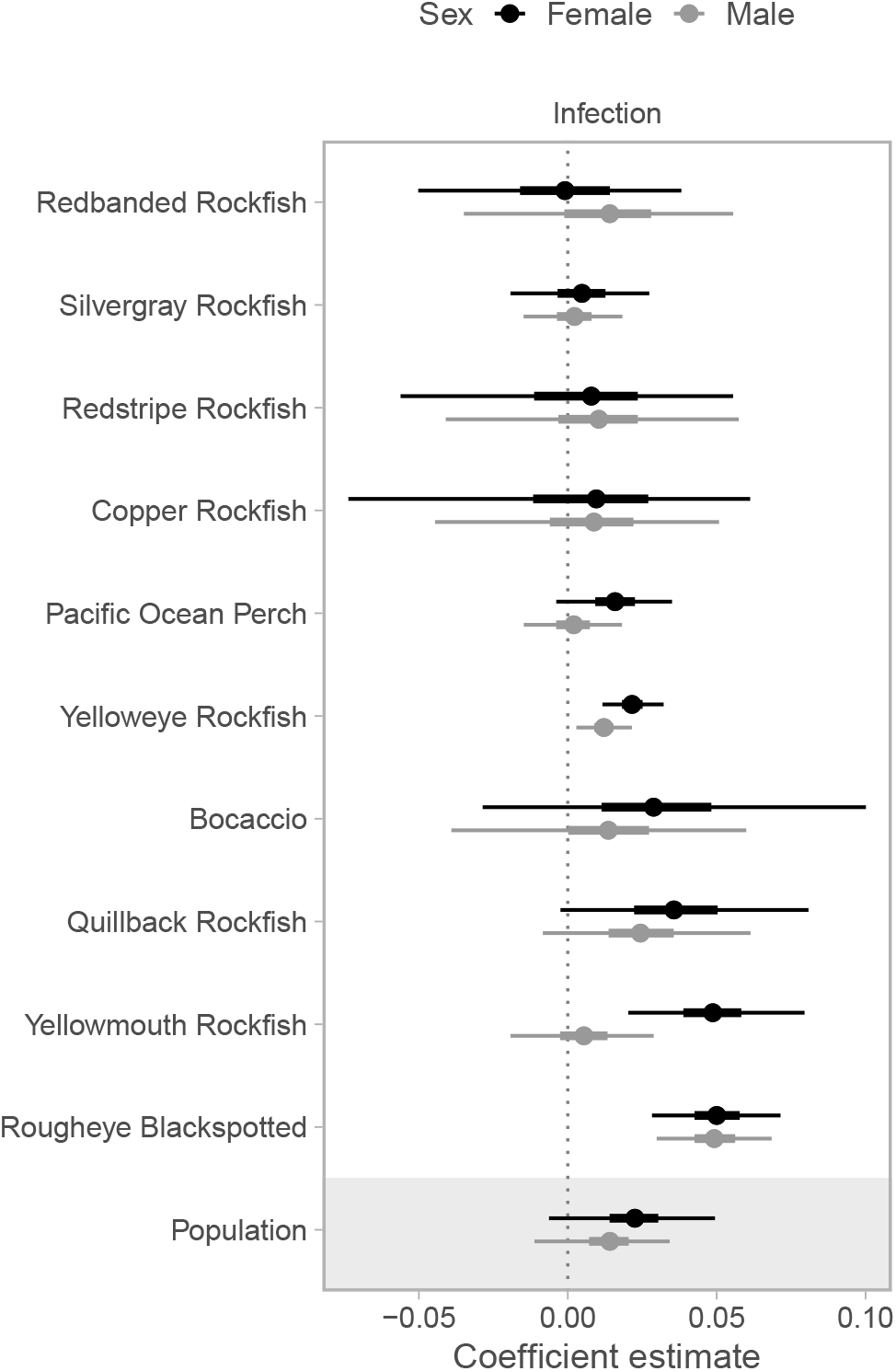
Posterior distribution of effect of *Sarcotaces* sp. infection on Le Cren’s body condition index (*K*). Values greater than 0 (vertical dashed line) indicate greater *K* in infected individuals than uninfected individuals and species are ordered by increasing effect of infection on *K*, with population-level differences shown at the bottom. Points are posterior medians and thick and thin lines show 50% and 95% credible intervals, respectively.

## 4 Discussion

To our knowledge, this is the first study to report on the prevalence of *Sarcotaces* sp. infection across many rockfish species from a large geographic area. We have observations from over 37,000 individuals that span 20+ years (and included 33 positive observations from 1969 and 1982). *Sarcotaces* sp. have been observed in fish caught across the entire survey frames for our Synoptic Trawl and Hardbottom Longline surveys, and at all depths that fish are captured. Infection rates are mostly below 2% for the majority of rockfish and thornyhead species encountered in our research surveys. However, a few species that have commercial and cultural importance (POP, Silvergray and Yelloweye Rockfish) exhibited infection rates that range from 2.9–10%. Infection rates were also found to be higher in immature fish relative to mature fish. We found that infection affects both male and female length-at-maturity ogives; however, the effect is more pronounced in males. For the three species that we generated a collective age-at-maturity ogive for, infection had a larger and negative effect on females, and the effect was weaker in males. Body condition was slightly higher in REBS, Yelloweye, and Yellowmouth with *Sarcotaces* sp. infection; however, it is likely that, from a biological perspective, the overall effect of infection on body condition was negligible. On the other hand, our results demonstrate a negative effect of *Sarcotaces* sp. infection on rockfish maturity ogives, and while preliminary, may affect other life-history traits as well.

Our research surveys do not effectively sample all rockfish species that occur in British Columbia coastal waters. Specifically, we do not catch Black, Tiger, or China rockfishes in any appreciable quantity, despite these species being caught more frequently in recreational fisheries (DFO Recreational Catch Statistics). Our understanding of infection rates within species; therefore, is biased by our catch rates of species encountered in our surveys. We mitigated this bias in developing the maturity ogives by including species as a random effect in our hierarchical model, which allowed us to estimate the effect for species with high catch rates while improving estimates for those with limited data. Future work that includes biological samples of other species more frequently caught in recreational fisheries would increase our understanding of infection across species. However, obtaining sufficient biological data from recreational fishers is logistically challenging.

Host-parasite interactions in the oceans are likely changing due to climate change (Harvell et al. 2002). Increasing water temperature is likely to be an important driver of changes in host-parasite interactions. These changes may be particularly relevant in temperate regions, where rising water temperatures—including both warmer summer highs and warmer winter lows (Cheng et al. 2021)—are occurring alongside extreme heat anomalies (Peterson et al. 2015, Amaya et al. 2020). Parasite transmission occurs primarily during the warm summer period (Karvonen et al. 2010), whereas parasite growth and recruitment is generally limited by cold temperatures during winter (Hakalahti et al. 2006). Rising temperatures may also increase parasite virulence by causing phenological mismatches, where infection occurs in the host during younger and more vulnerable life stages (Paull and Johnson 2011).

Rockfish densities are, concurrently, also expected to change with increasing ocean temperatures (English et al. 2021). English et al. (2021) modelled local changes in biomass density for 38 demersal fishes in British Columbia. In localities with relatively cooler temperatures, positive biotic velocities (i.e., increasing biomass) were associated with increasing temperatures, while in relatively warmer localities, negative biotic velocities (i.e., decreasing biomass) were associated with additional warming. Interestingly, they found that Yelloweye Rockfish was most likely to experience local density declines with increasing temperature, which is the species we found to have the highest rate of *Sarcotaces* sp. infection (but did not find an effect on maturity). Scant baseline information on parasite prevalence, virulence, and effects on life history for all rockfish species, however, makes it challenging to understand how these interactions are changing, and what the consequences are for populations and communities.

Many of the rockfish species examined in this study are socio-economically important. They are targeted in commercial, recreational, and First Nations’ Food, Social, and Ceremonial fisheries. Stock assessments estimate key population parameters such as growth, natural mortality, maturity, biomass, and recruitment to understand stock productivity and provide catch advice to fisheries managers. Fish health, despite the obvious link between health and productivity, is not routinely considered in stock assessments (Lloret et al. 2012); the effect of parasites on fish life history are considered even less frequently. Considering the dearth of data we have on fish host-parasite interactions, this is unsurprising. However, this does not negate a possible negative health effect of *Sarcotaces* sp. infection on rockfish.

Our analysis of infection prevalence across maturity stages shows a greater proportion of infected immature fish relative to fish that are reproductively mature. Likewise, our age data, although limited, also shows that the mean age of infected individuals is lower than that for uninfected individuals. We do not have sufficient data to conclude that infection increases mortality before individuals reach reproductive maturity; however, our results show this is a hypothesis worth further investigation.

It is difficult to know how the *Sarcotaces* sp. infection rates we observed in British Columbia rockfishes compares to the infection rates in other rockfish populations, other groundfishes (aside from *Antimora microlepis*), and the infection rates of other copepod and non-copepod parasites. For such commercially and socially important species as POP and Yelloweye Rockfish, infection rates of 2.9% and 10%, respectively, may be sufficiently high to warrant follow-up studies. Expanding biological sampling to include fecundity would permit exploring the effect of infection on an additional key life-history trait, as was suggested for Silvergray Rockfish by Stanley and Kronlund (2005). Additional condition indices, such as the hepatosomatic index (liver weight / body weight) may be more applicable for detecting the physiological cost of parasitism than condition indices based on weight-length relationships (Hasegawa and Poulin 2025). Finally, testing the sensitivity of key stock assessment parameters to different maturity and fecundity values would help determine whether, and how, accounting for *Sarcotaces* sp. infection affects catch advice.

## Acknowledgments

The authors thank the Groundfish Surveys Team, both past and present, for their leadership and efforts on all the groundfish research surveys, and the Captains, Fishing Masters, and crews of the CCGS Fishery Research Vessels and industry partners for their participation in the groundfish research surveys. The authors also thank the DFO Sclerochronology Lab for their efforts processing thousands of otoliths a year for age data.

## Data availability

Data and code used in this study are available at https://github.com/pbs-assess/sarc and will be archived on Zenodo prior to publication. Additionally, all DFO Groundfish survey data is available on Open Canada (open.canada.ca)

