## Supporting Information for "Prevalence of the parasitic copepod, *Sarcotaces* sp., infection in British Columbia rockfishes (*Sebastes* sp.) and implications for rockfish life-history"

### Detailed Methods

#### Spatial modelling of *Sarcotaces* sp. infection probability

We modeled the spatial probability of *Sarcotaces* sp. encounter with a logistic regression with a quadratic effect for log depth, an independent mean for each year, and a Gaussian Markov random field (GMRF) to account for latent spatial factors driving spatial correlation. Specifically our model was of the form:

$$y_i^{\text{sarc}} \sim \text{Bernoulli}(\pi_i^{\text{sarc}}), \tag{S1}$$

where the probability of encountering an infected fish, $\pi_i^{\text{sarc}}$, is modeled as:

$$\text{logit}\left(\pi_i^{\text{sarc}}\right) = \alpha_t + \beta_1 D_i + \beta_2 D_i^2 + \omega_{\boldsymbol{s}}. \tag{S2}$$

Here, $y_i^{\text{sarc}}$ is an observation for fish $i$ that is either (1) infected with *Sarcotaces* sp. or (0) not infected with *Sarcotaces* sp., $\alpha_t$ is a year-specific mean, $\beta_1$ and $\beta_2$ are coefficients on the effects of log depth ($D_i$) and log depth squared ($D_i^2$) for fish $i$, and $\omega_{\boldsymbol{s}}$ is a value from a spatial GMRF representing the spatial effect of latent variables at spatial coordinates $\boldsymbol{s}$. The vector of $\omega_{\boldsymbol{s}}$ values, $\boldsymbol{\omega}$, is assumed drawn from a multivariate normal distribution with covariance (inverse precision) matrix $\boldsymbol{\Sigma}_{\boldsymbol{\omega}}$:

$$\boldsymbol{\omega} \sim \text{MVN}\left(\mathbf{0}, \boldsymbol{\Sigma}_{\boldsymbol{\omega}}\right). \tag{S3}$$

We allowed for anisotropy in the GMRF (Haskard 2007, Fuglstad et al. 2015) to allow for spatial correlation to decay differently up and down the coast from off the coast and fit the model using a finite-element mesh (Lindgren et al. 2011, Lindgren 2025) with a minimum triangle edge length of 5 km. We fit this spatial model with the sdmTMB R package (Anderson et al. 2024), which uses fmesher (Lindgren 2025) to generate input matrices and TMB (Kristensen et al. 2016) to calculate the marginal likelihood.

#### Bayesian modelling of length- and age-at-maturity

We estimated length-at-maturity with hierarchical Bayesian logistic regression models fit separately to male and female fish:

$$y_i^{\text{mat}} \sim \text{Bernoulli}(\pi_i^{\text{mat}}), \tag{S4}$$

where the probability of maturity, $\pi_i^{\text{mat}}$, is modeled as:

$$\operatorname{logit}\left(\pi_i^{\text{mat}}\right) = \beta_0 + \beta_1 L_i + \beta_2 I_i + \beta_3 L_i I_i \\ + b_{0,s[i]} + b_{1,s[i]} L_i + b_{2,s[i]} I_i + b_{3,s[i]} L_i I_i. \tag{S5}$$

Here, $y_i^{\text{mat}}$ is either mature (1) or immature (0) for fish $i$; $\beta_0$, $\beta_1$, $\beta_2$, and $\beta_3$ are fixed-effect coefficients; $L_i$ is the (standardized) length of fish $i$; $I_i$ is the presence (1) or absence (0) of *Sarcotaces* sp. infection in fish $i$, and $b_{0,s[i]}$, $b_{1,s[i]}$, $b_{2,s[i]}$, and $b_{3,s[i]}$ are species-specific random intercepts and slopes for species $s$, fish $i$. The vector of species-specific random effects $\boldsymbol{b}$ $(b_0, b_1, b_2, b_3)$ are modeled as:

$$\boldsymbol{b} \sim \text{MVN}\left(\boldsymbol{0}, \boldsymbol{\Sigma_b}\right), \tag{S6}$$

where $\boldsymbol{\Sigma_b}$ is a covariance matrix of the variances and covariances between the random intercepts and slopes.

We assigned Gaussian priors on all $\beta$ coefficients:

$$\beta_0 \sim \text{N}(0, 10^2), \tag{S7}$$

$$\beta_1, \beta_2, \beta_3 \sim \text{N}(0, 2^2), \tag{S8}$$

a half-Student-t prior with a degrees of freedom 3 on the standard deviations of all random effects:

$$\sigma_0, \sigma_1, \sigma_2, \sigma_3 \sim \text{Half-t}(3, 0, 2), \tag{S9}$$

and an LKJ prior on the correlation matrix, $\boldsymbol{R}$, of the random effects:

$$\boldsymbol{R} \sim \text{LKJ}(1), \tag{S10}$$

where $\boldsymbol{\Sigma_b}$ is formed as:

$$\boldsymbol{\Sigma_b} = \boldsymbol{DRD}, \tag{S11}$$

where $\boldsymbol{D}$ is the diagonal matrix of standard deviations from $\boldsymbol{\Sigma_b}$.

Using the notation $[\boldsymbol{y}|\boldsymbol{x}]$ to denote the distribution of $\boldsymbol{y}$ conditional on $\boldsymbol{x}$, we can write the proportional joint probability density as:

$$[\boldsymbol{\beta}, \boldsymbol{b}, \boldsymbol{\Sigma_b} \mid \boldsymbol{y}] \propto [\boldsymbol{y} \mid \boldsymbol{\beta}, \boldsymbol{b}, \boldsymbol{\Sigma_b}]\,[\boldsymbol{b} \mid \boldsymbol{\Sigma_b}]\,[\boldsymbol{\Sigma_b}]\,[\boldsymbol{\beta}] \tag{S12}$$

Since there were fewer data with ages available (total n = 2,334, infected n = 37), we were unable to fit the above hierarchical model to estimate age-at-maturity. Instead, we fit a simplified model of the probability of age-at-maturity as:

$$\text{logit}\left(\pi_i^{\text{mat}}\right) = \beta_0 + \beta_1 A_i + \beta_2 I_i + \beta_3 A_i I_i \tag{S13}$$

where symbols are as described above with the addition of $A_i$ representing standardized age of fish $i$. We used the same priors as in Equations S7 and S8.

To compare *Sarcotaces* sp. infection probability between fishes of different sex and maturity status levels (Appendix 1), we fit a separate hierarchical logistic regression of the form:

$$\text{logit}\left(\pi_i^{\text{mat}}\right) = \boldsymbol{X}_i \boldsymbol{\beta} + \boldsymbol{X}_i \boldsymbol{b}_{s[i]} \tag{S14}$$

where $\boldsymbol{X}_i$ is a vector of predictors from a model matrix of the interaction between maturity bins and sex, $\boldsymbol{\beta}$ is a vector of associated fixed effect coefficients, and $\boldsymbol{b}_{s[i]}$ is a vector of associated random effects. The random effects were modelled as before:

$$\boldsymbol{b} \sim \text{MVN}\left(\boldsymbol{0}, \boldsymbol{\Sigma}_{\boldsymbol{b}}\right), \tag{S15}$$

and the same priors were used as in Equations S7–S11.

### Bayesian modelling of body condition

For body condition, we fit a hierarchical model as described in the main text. We used the same MVN random effect distribution and priors as described in equations S6–S11. The proportional density can be described as in equation S12.

### Bayesian modelling of age as a function *Sarcotaces* sp. infection

We modelled age as a function of infection status with a gamma generalized linear model with a log link. We applied the same $\text{N}(0, 2^2)$ and $\text{N}(0, 10^2)$ priors to the slope and intercept parameters and a $\text{gamma}(0.1, 0.1)$ prior to the shape parameter.

### Supporting Tables

Table S1: Rockfish maturity stages and descriptions. Stages 1 and 2 are classified as immature, stages 3-7 as mature.

| Sex | Maturity Code | Stage Name | Description |
|---|---|---|---|
| Female | 1 | Immature | Ovaries are small, translucent, no visible eggs |
| | 2 | Maturing | Ovaries small but swelling, semi-translucent, no visible eggs, no black dots, granular |
| | 3 | Developing | Ovaries medium sized, visible eggs, eggs opaque |
| | 4 | Fertilized | Ovaries large, hydrated eggs, less than half of eggs translucent |
| | 5 | Running Ripe | Ovaries large, eyed larvae, larvae translucent with black eyes |
| | 6 | Spent | Ovaries large and flaccid, red in color, few leftover eggs present |
| | 7 | Resting | Ovaries medium size with firm, thicker membrane, splotchy grey in color |
| Male | 1 | Immature | Testes small and string-like, translucent in color |
| | 2 | Maturing | Testes swelling, translucent in color |
| | 3 | Developing | Testes medium in size, swelling, beige-whitening in color |
| | 4 | Developed | Testes large and swollen, easily broken, white in color |
| | 5 | Running Ripe | Testes large and swollen with running sperm, white in color |
| | 6 | Spent | Testes large and firm with thick milt, tan and white color in cross section |
| | 7 | Resting | Testes medium in size, triangular in cross section, brown in color |

Table S2: Frequency of the number of *Sarcotaces* sp. cysts found within each individual combined across species

| # cysts | 1 | 2 | 3 | 4 | 5 | 6 | 7 | 8 | 9 | 10 |
|---|---|---|---|---|---|---|---|---|---|---|
| N | 35,281 | 976 | 144 | 38 | 6 | 8 | 2 | 5 | 3 | 1 |

Table S3: Sample sizes by maturity status, infection status, and sex. For the analysis, maturity stages 4 and 5 were combined due to low sample sizes.

| | | Female | | Male | |
|---|---|---|---|---|---|
| Maturity | Maturity status | Uninfected | Infected | Uninfected | Infected |
| Immature | 1 | 901 | 83 | 1,363 | 157 |
| | 2 | 2,625 | 142 | 2,525 | 178 |
| Mature | 3 | 2,530 | 52 | 4,502 | 117 |
| | 4 | 259 | 14 | 1,362 | 15 |
| | 5 | 319 | 7 | 229 | 0 |
| | 6 | 1,911 | 49 | 1,357 | 37 |
| | 7 | 5,689 | 200 | 5,118 | 118 |

### Supporting Figures

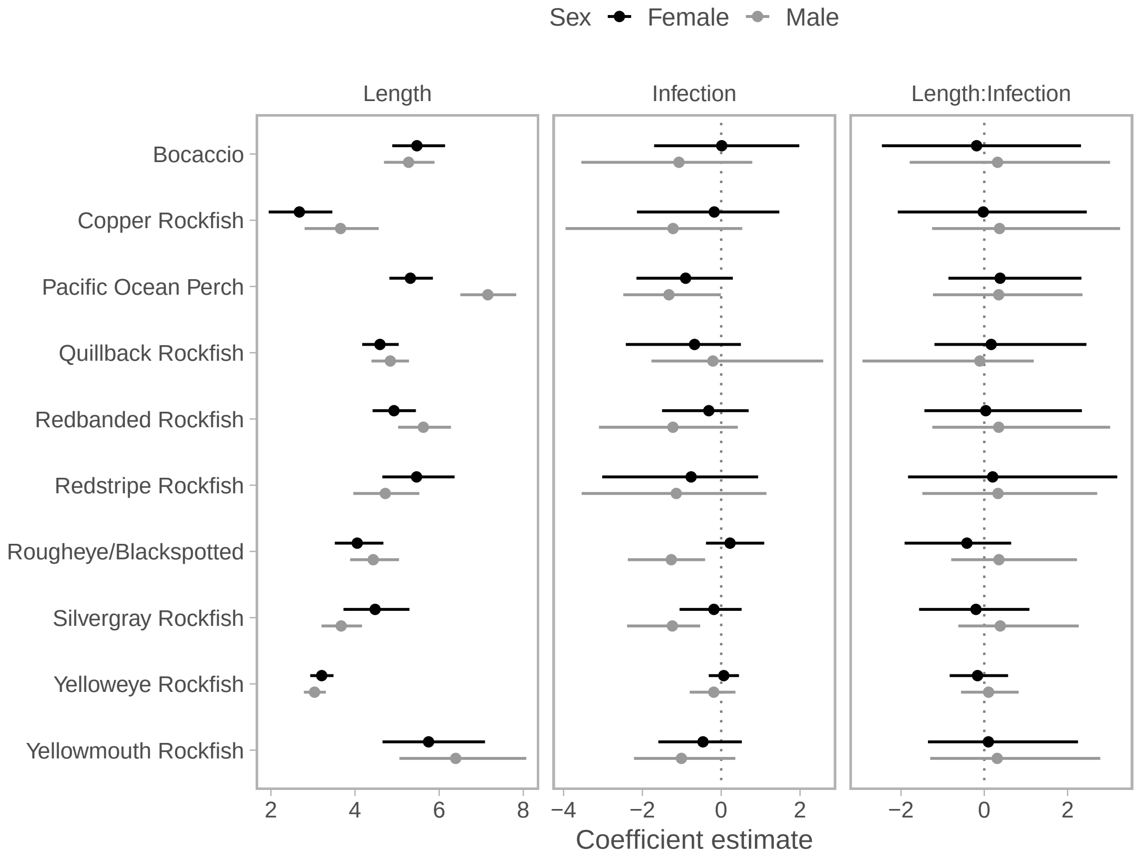


Figure S1: Coefficient estimates from the hierarchical model of fish length and infection on maturity. Dots are posterior medians and line segments are 95% credible intervals.

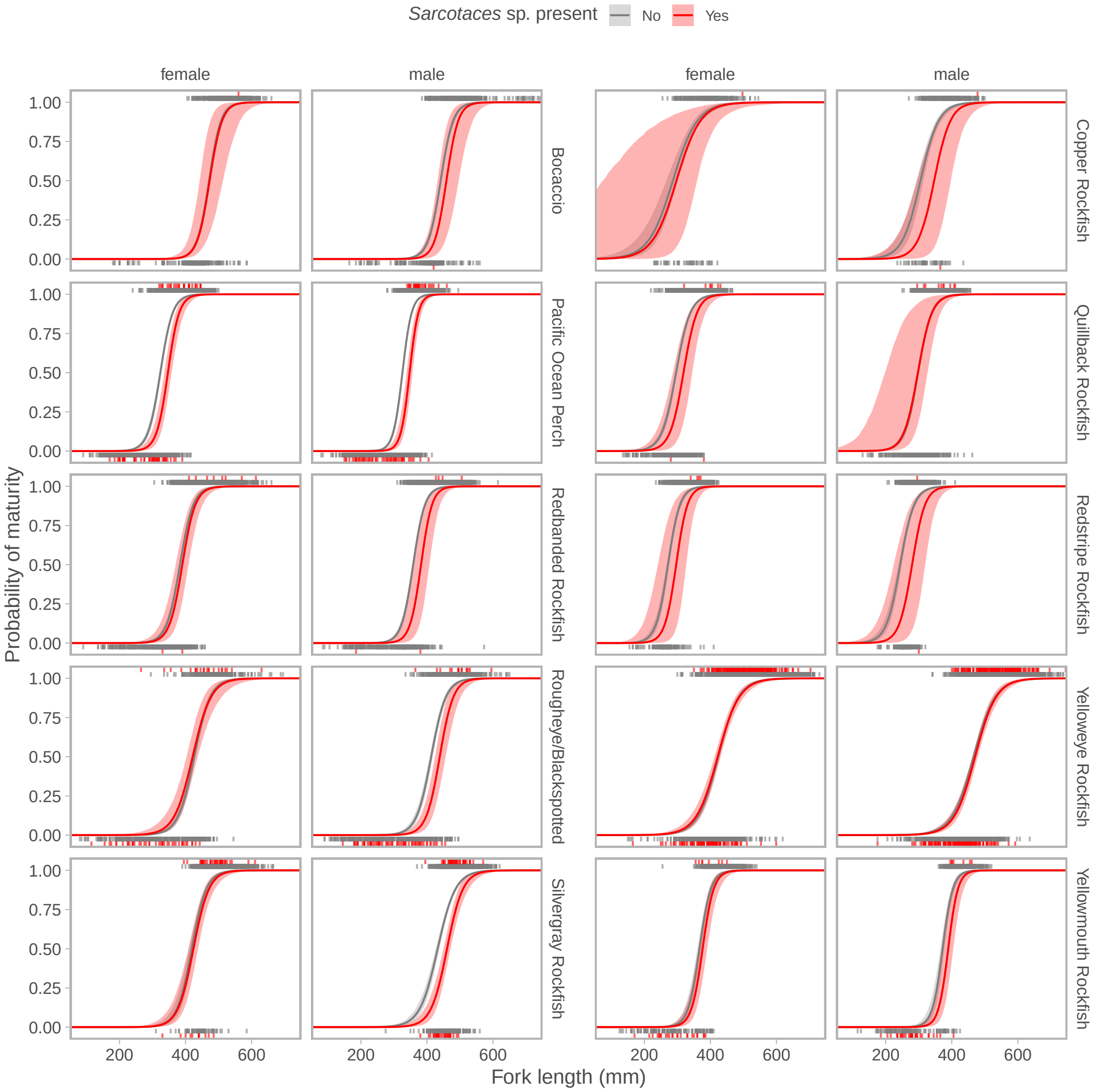


Figure S2: Length at maturity ogives for select rockfish species (>10 infected individuals) separated by infection status and by sex. Shaded areas represent 95% credible intervals and lines represent medians.

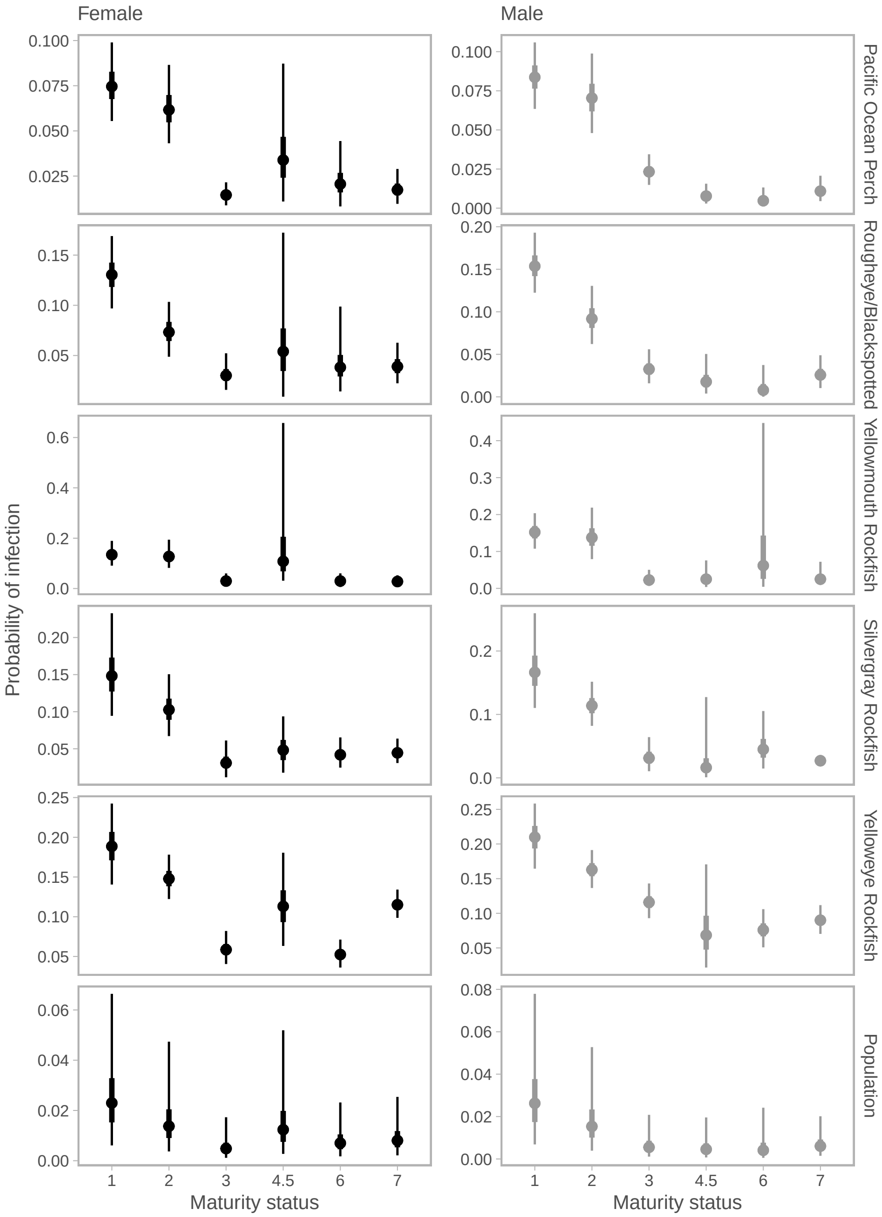


Figure S3: Probability of infection in female and male fish by maturity code. Line segments represent 95% and 50% credible intervals and dots represent medians. Species are ordered from top to bottom in increasing infection rates of maturity status 1 (status: immature), with the population-level infection rates shown at the bottom. Only species with at least 10 infections found in each of immature (status 1 and 2), and mature (status 3 to 7) are shown.

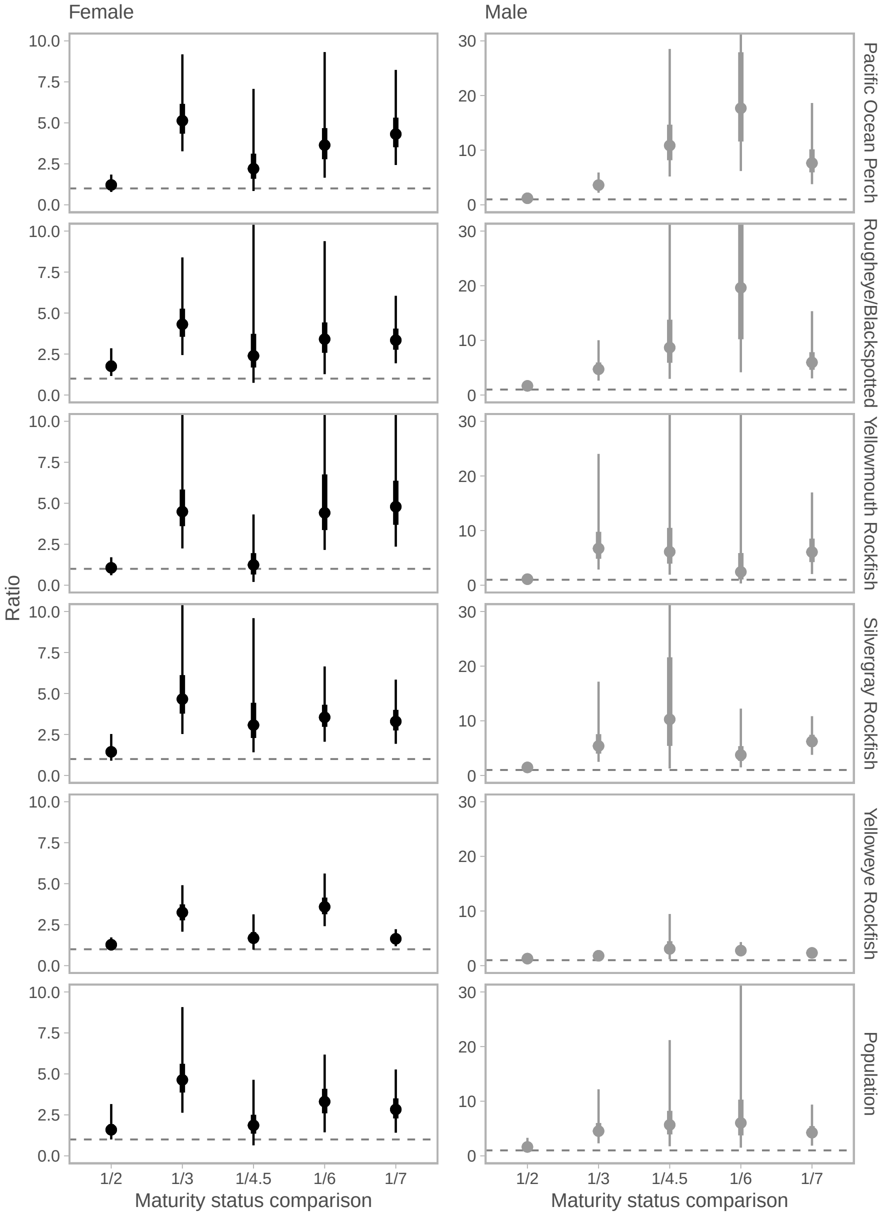


Figure S4: Posterior distribution of the ratio of infection probabilities in maturity status 1 (immature) compared to all later maturity statuses. The ratio is calculated as Pr(infection|status 1) / Pr(infection|status > 1). Values greater than 1 (vertical dashed line) indicate higher infection probability in immature individuals. Points are posterior medians and thick and thin lines show 50% and 95% credible intervals, respectively. Species are ordered from top to bottom by increasing ratio of infection rate, with the population-level difference shown bottom.

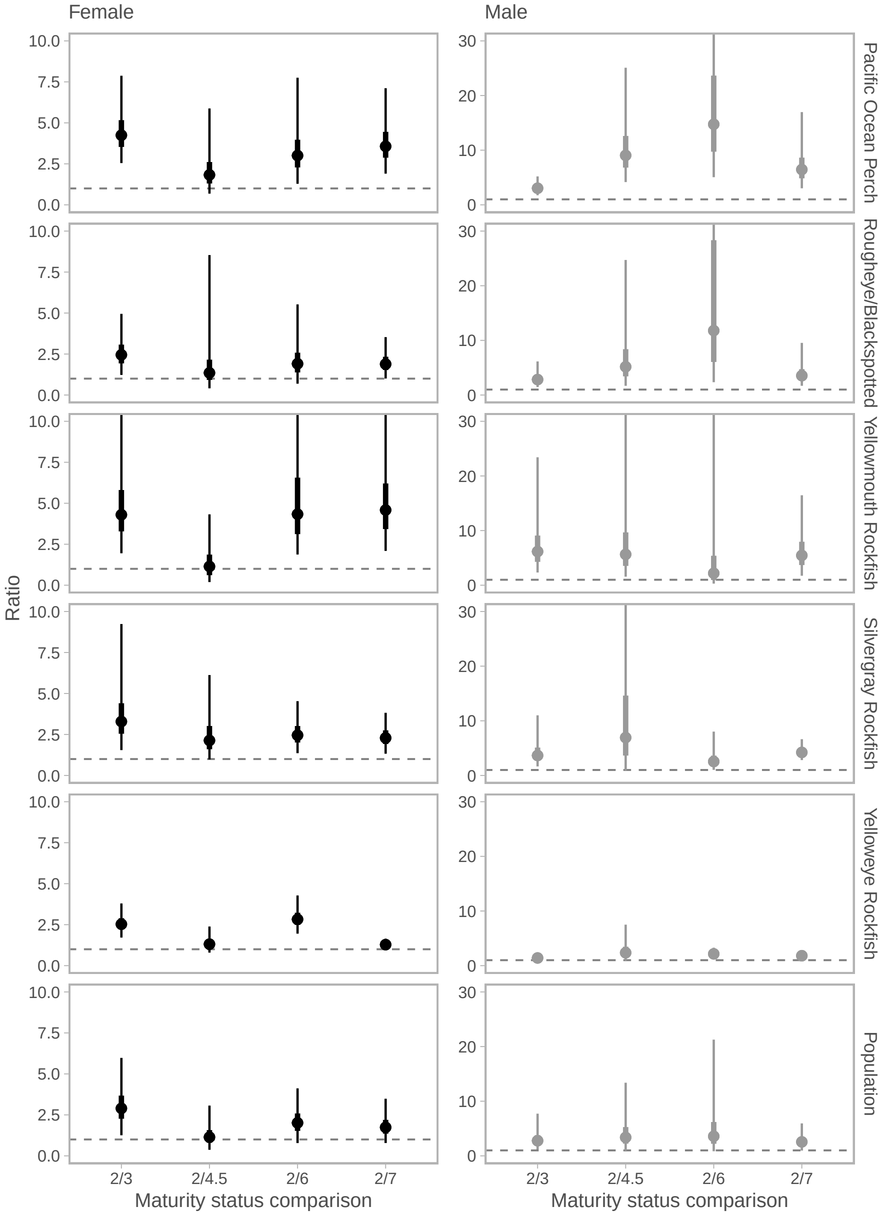


Figure S5: Posterior distribution of the ratio of infection probabilities in maturity status 2 (maturing) compared to all later maturity statuses. The ratio is calculated as Pr(infection|immature) / Pr(infection|mature). Values greater than 1 (vertical dashed line) indicate higher infection probability in immature individuals. Points are posterior medians and thick and thin lines show 50% and 95% credible intervals, respectively. Species are ordered from top to bottom by increasing ratio of infection rate, with the population-level difference shown bottom.

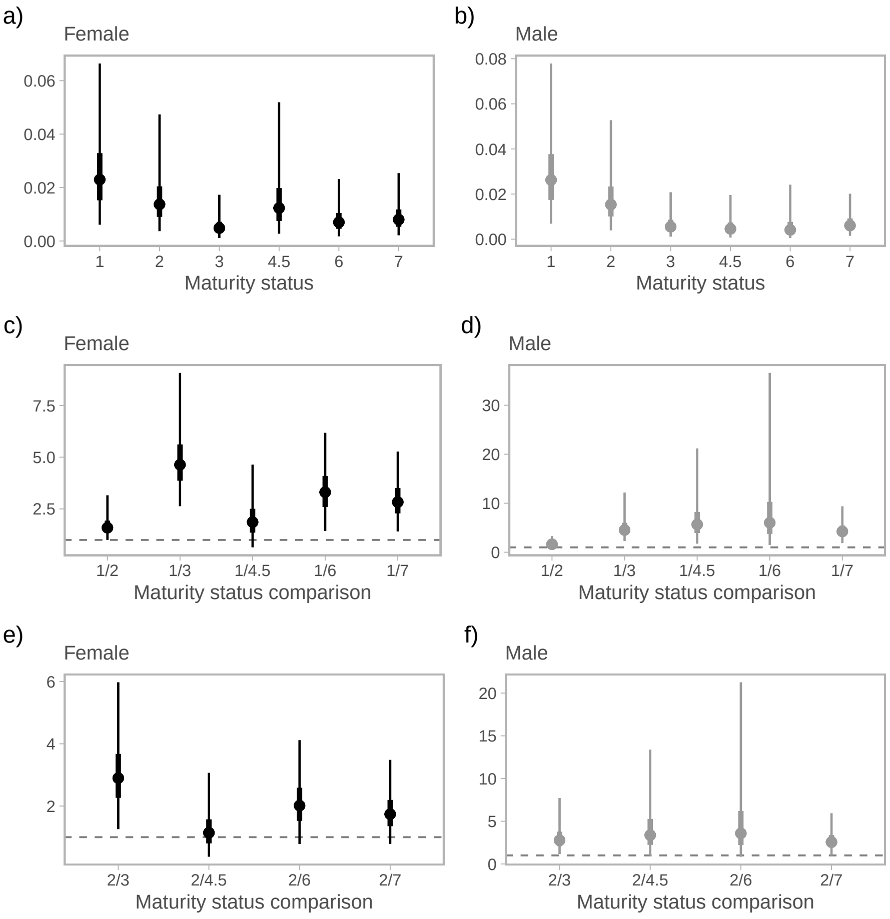


Figure S6: Population-level infection probability patterns across fish maturity stages by sex. (a,b) Posterior distributions of infection probability in female and male fish across maturity codes 1–7, where codes 1–2 are reproductively immature fish (1 = immature, 2 = maturing) and codes 3–7 are reproductively mature fish. (c,d) Posterior distributions of infection probability ratios comparing immature fish (maturity code 1) to each later stage individually (codes 2, 3, 4.5, 6, 7), calculated as (Pr(infection|code 1) / Pr(infection|code $i$) for each code $i$. (e,f) Posterior distributions of infection probability ratios comparing maturing fish (maturity code 2) to each mature stage individually (codes, 3, 4.5, 6, 7), calculated as (Pr(infection|code 2) / Pr(infection|code $i$) for each code $i$. Maturity codes 4 and 5 were combined due to small sample sizes (S3). In panels c–f, ratios greater than 1 (horizontal dashed line) indicate higher infection rates in the younger maturity group. Points are posterior medians and thick lines show 50% and thing lines show 95% credible intervals.
